# Automated behavioral segmentation and markerless pose tracking of mice during spaceflight

**DOI:** 10.64898/2026.04.30.721950

**Authors:** Frederico C. Kiffer, Ryan T. Scott, Marlu T. Martens, Arthur Mayo, Yipeng Li, Mariela Mendoza, Shambhabi Gautam, Julian Huang, Meera Bathwal, Svanik Jaikumar, Amishi Mahajan, Lauren M. Sanders, Amelia J. Eisch, Talmo D. Pereira

**Author notes:** Co-corresponding authors: Frederico Kiffer, Amelia Eisch, Talmo Pereira.

## Abstract

The NASA Rodent Habitat aboard the International Space Station enabled long-duration studies of behavioral responses to spaceflight, but video-based behavioral analysis has relied on laborious manual annotation. No study has tested whether deep learning tools can automate this analysis under the demanding imaging conditions of orbital vivaria. We applied pose estimation (SLEAP) and behavioral segmentation (DeepEthogram) to archival footage from the Rodent Research-1 mission. Nine labelers annotated 3,249 pose labels across 2,063 frames, and three behaviorists labeled 411,194 frames across 66 videos. Pose tracking accuracy approximated human inter-annotator variability despite progressive lens soiling, grid occlusions, and spherical aberration. Behavioral classification across eight categories achieved accuracy of 0.86-0.90 and suggests progressive behavioral adaptations to microgravity. Kinematic reconstruction of circling estimated centripetal accelerations periodically approaching 1*g*. This is the first application of deep learning-based pose estimation and behavioral segmentation to rodents in spaceflight, establishing benchmarks for future monitoring systems.

**One-Sentence Summary:** Deep learning-based pose estimation and behavioral segmentation of ISS video reveal progressive behavioral adaptations of mice to microgravity

---

Enabling long-duration and long-distance human spaceflight requires understanding spaceflight risk factors such as microgravity, ionizing radiation, hypercapnia, regolith exposure, and isolation-confinement (*1*). Ground analog models exist to replicate certain spaceflight conditions but remain imperfect, making studies of model organisms in actual spaceflight or interplanetary missions critical for understanding human physiological responses and systems biology, and for developing countermeasures. Mice have served as a spaceflight model organism for decades, with use through the Apollo era for in-flight radiobiology (*2*) and throughout the Space Shuttle program, where neurological and behavioral effects (*3*, *4*) were among the key research foci until its final mission in 2011 (*5*). Mouse housing aboard the International Space Station (ISS) began in 2009 with the Italian Space Agency’s Mouse Drawer System, which was limited to single housing, constraining sample size and inducing social isolation stress (*6*). In 2014, NASA reintroduced mouse spaceflight research with the Rodent Research-1 (RR-1) mission, using the Animal Enclosure Module–eXtended (AEM-X) vivarium housed in an EXPRESS rack aboard the ISS (*7*). This system contains a central food block, pressurized water activated by Lixits, and four analog cameras used throughout the mission for research and veterinary evaluation of gross rodent health. Subsequent ISS missions also saw the Japan Aerospace Exploration Agency (JAXA) deploy their Mouse Habitat Unit in the Kibo module, a centrifuge-based vivarium capable of replicating terrestrial, lunar, Martian, and microgravity conditions (*8*).

Microgravity produces a cephalic fluid shift that is hypothesized to contribute to Spaceflight-Associated Neuro-ocular Syndrome (SANS), a set of neuro-ophthalmic findings whose underlying pathophysiology remains under investigation (*9*). In spaceflight analog studies combining head-down tilt bed rest with ISS-level CO₂, hypercapnia reduced cerebral perfusion (*10*), and altered brain connectivity and sensorimotor performance (*11*). CO₂ levels aboard the ISS correlate with headaches experienced by crew members (*12*). Although the ISS is well within Earth’s radioprotective magnetosphere, astronauts still experience some space radiation exposure, which is known to alter rodent behavior at various dosages (*13*). Long-duration spaceflight also appears to result in structural brain changes in astronauts, including evidence of narrowing of the central sulcus, upward shift of the brain, and ventricular enlargement that exceeds normal aging (*14*, *15*). Astronauts show hippocampal volumetric changes following ISS missions (*16*), and an RR-1 study resulting from the same mice in the present work suggests that spaceflight decreases inflammasome signaling in the brain when compared to ground-matched controls and pre-flight baseline readings (*17*). Behavioral health and performance decrements rank among the highest-priority risks for exploration-class missions (*18*). Isolation, confinement, and psychosocial stressors further compound spaceflight-induced neurological changes, affecting crew mood, cognition, and interpersonal dynamics (*19*, *20*). Given the breadth of these spaceflight-induced changes to the central nervous system (CNS) (*21–24*), behavioral monitoring becomes essential both as a readout of CNS function and as a practical tool for assessing crew and subject well-being during missions where real-time ground support is limited. Rodent models provide a tractable system for developing and validating such monitoring and countermeasure approaches before they can be adapted for crew. However, current behavioral analysis of rodents in spaceflight relies on manual scoring, a process that is labor-intensive, difficult to standardize, and poorly suited to the scale of longitudinal monitoring required for exploration class missions.

To these ends, researchers at NASA Ames retrospectively analyzed archival video footage of RR-1, for indices of rodent behaviors and activity (*25*). The RR videos were collected originally as part of the mission ‘health check’ videos to monitor subject well-being aboard the ISS, which was an Institutional Animal Care and Use Committee (IACUC) requirement. The researchers manually scored videos for a limited number of behaviors, and occlusion analyses were performed to measure camera view soiling across mission days. Although some attempts were made to compare the behaviors of mice in flight to ground behaviors, these efforts were frustrated by uneven sampling across cages, time, and camera location. However, this laborious analysis found that lens soiling significantly occluded the field of view from experimental day 15 onward, and that mice showed locomotor adaptations involving the shift from forelimb to quadrupedal locomotion over time, and the emergence of a circling behavior, in which mice ran around the four walls of the cage in a running wheel-like fashion, likely loading their musculoskeletal system (*25*).

Since this work was performed, open-source computer vision tools for high-accuracy visual tracking with common computational hardware have proliferated, including SLEAP (Social LEAP Estimates Animal Poses), a pose estimation tool developed by our group (*26*). Recent work has used these types of approaches to understand how plants, *C. elegans*, *D. melanogaster*, and *D. rerio* adapt to spaceflight (*27*). One group captured mouse activity in the JAXA Habitat Cage Unit of the Kibo module using conventional, non-deep-learning tracking tools (*28*). However, no work has applied and evaluated the technological readiness level of deep learning tools to capture and study rodent pose and behavior in space. Rodent pose and behavior are challenging to capture in space due to numerous imaging obstacles, including spherical aberration, varying video resolution, complex 3-dimensional views, inability to distinguish individuals, and progressive soiling of the camera lens (*29*).

To assess the feasibility of automating longitudinal kinematic and behavioral capture of freely behaving mice in space, we accessed archival footage of RR-1 from the NASA Open Science Data Repository (OSDR), specifically from the NASA Ames Life Sciences Data Archive (ALSDA) (*30*, *31*), and curated and standardized videos across the mission. We then applied the deep learning-based pose estimation tool SLEAP and the behavioral segmentation tool DeepEthogram to this dataset. Here, we benchmark automated pose tracking against human annotators, classify eight behavioral categories across the mission, apply kinematic reconstruction to estimate centripetal forces generated during circling behavior, and share labeled data, trained models and predicted results. We hope this work serves as a basis with which to inform future system design and establish a baseline for technical readiness level of technology for automating behavioral quantification in future space biology research studies.

## Results

### Video Sampling

The RR-1 video dataset consists of videos sampled throughout a 37-day mission and contains footage from the two Lixit cameras and two Filter cameras on each side of the AEM-X vivarium. Lixit camera views are extremely close to the mice, and offer a very limited field of view, making them unsuitable for behavioral analysis. Therefore, only footage from the Filter cameras was used. The original dataset consisted of video uploads from the ISS that were originally intended to include 30 min of footage at the start of each light cycle daily, but a thermal runaway issue instead led to a wide distribution of video resolutions among ground controls and flight cages (**Fig. S2a**). Flight videos had a somewhat more uniform distribution of video resolutions. All dark cycle flight videos were 640 x 480 pixels, and light cycle videos were either the same resolution, higher resolution, or slightly below 640 x 480 pixels (**Fig. S2b**). Therefore, we standardized all flight videos to 640 x 480 pixels. Issues related to sampling also meant that videos were captured at varying lengths, with non-Gaussian distributions (Light: *P*<0.0001; Dark: *P*<0.0001; **Fig. S2c**), and non-Gaussian distributions of sampling days with respect to light cycle (Light: *P*<0.0001; Dark: *P*<0.0001; **Fig. S2e**). We observed a significant difference in video lengths by light cycle, with light cycle videos containing on average 7,337 more frames (*P* < 0.0001; **Fig. S2d**). Day sampling disparity was also significant, with more sampling in the dark cycle (*P* < 0.0001; **Fig. S2f**). Furthermore, dark cycle flight videos were sampled near daily, whereas light cycle flight videos were sampled almost exclusively in the first half of the mission.

### Pose Estimation

To establish a human baseline for keypoint localization variation, nine labelers independently annotated three keypoints (snout, neck, and tail-torso interval [TTI]) across an identical dataset consisting of Filter 1 and 2 view frames, and containing diverse pose orientations and distances from the camera. Inter-annotator variability, measured as pixel distance from the consensus keypoint, differed by body part: the snout showed the lowest variability, followed by the neck and TTI (**Fig. 2a-d**). Variability also differed by subject position relative to the camera, with instances closer to the camera or near frame edges producing greater disagreement among labelers due to an inherently higher pixel size per keypoint (**Fig. 2a, b**). The nine annotators labeled a large representative dataset spanning all Filter cameras, light cycles, and the full mission duration to capture the progressive effect of lens soiling. This dataset was used to train a bottom-up SLEAP model with a snout-TTI and neck-TTI skeleton. The model performed well despite the challenges present in the video dataset. Raw localization errors averaged 20 or fewer pixels from the ground truth per keypoint, with the neck producing the largest average errors and the TTI the smallest (**Fig. 3b**). These distances are swayed by outliers, with most inferences showing error distances of less than 10 pixels (**Fig. 3c**). These uncalibrated error metrics remained anatomically relevant through the 75th percentile across all keypoints. However, the localization error at the 90th percentile was large (**Fig. 3a**). When model error was adjusted directly to human inter-annotator variability, model performance fell within range of human error, averaging 7 or fewer pixels from consensus/adjusted ground truth (**Fig. 3e**). Calibrated error distances at the 90th percentile remain close to ground consensus, suggesting that when calibrated for inter-annotator variability, the model approximates human-level localization accuracy (**Fig. 3d, f**).

**Figure 1.**
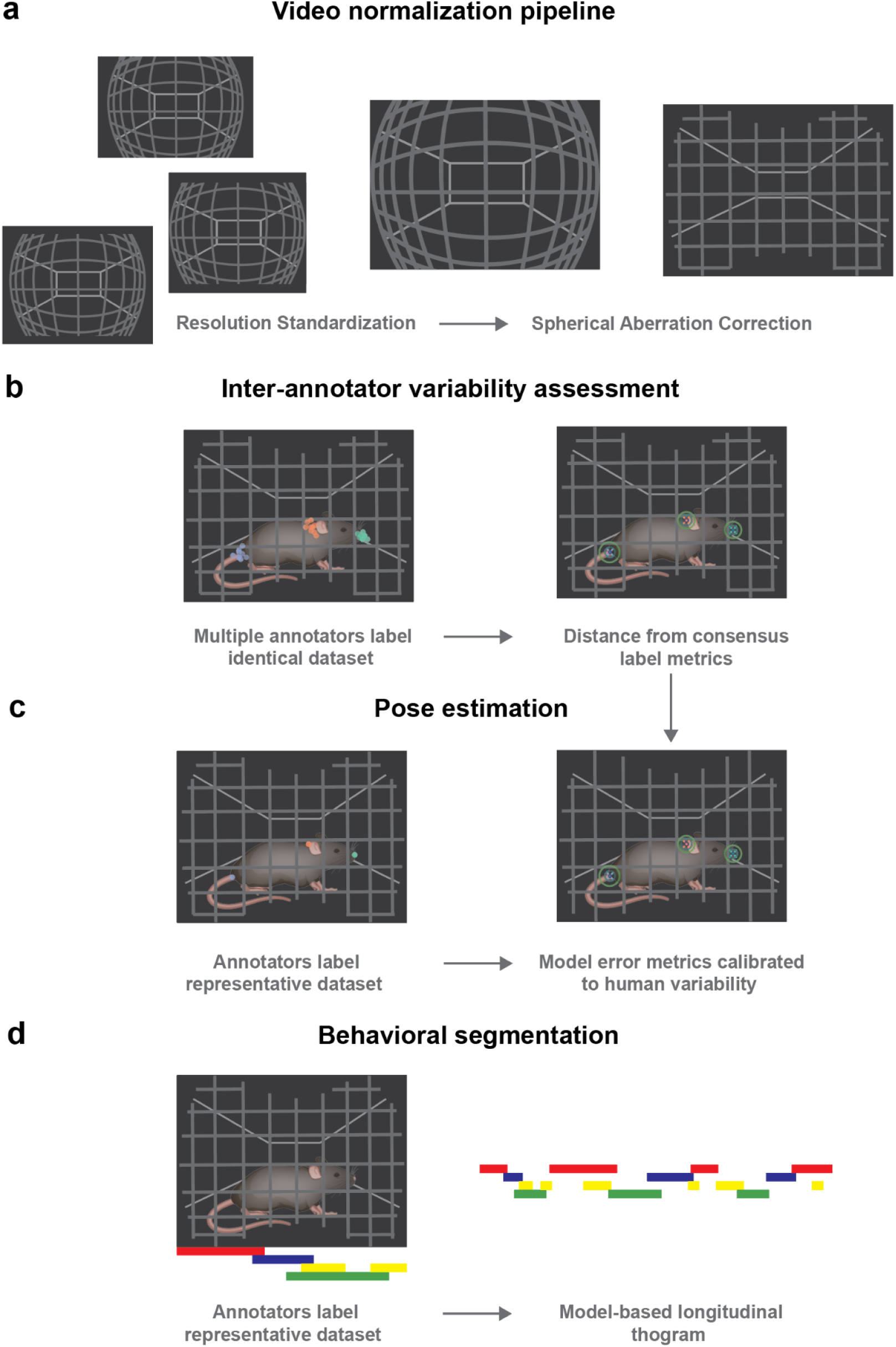
Experiment overview. **A.** Preprocessing pipeline for normalization of archived RR-1 videos. Videos are first aligned and standardized to a common resolution, then spherical aberration is corrected and standardized via intrinsic video properties. **B.** Pose annotation variability is assessed by 9 trained labelers on an identical dataset of 18 instances on raw videos. Labels are translated first to the video resolution standardization steps, then through the undistortion procedure. **C.** Pose estimation is performed with a ground truth dataset consisting of 3,249 labels by 9 trained annotators, and trained with the SLEAP pipeline on normalized videos. Model metrics are calibrated to human variability metrics. Kinematic analyses are performed on tracked inferences with smoothed interpolated points. **D.** A behavioral dataset consisting of 611,045 annotated behavioral event-frames across the mission duration was used to train the behavioral segmentation pipeline DeepEthogram, on normalized videos.

**Figure 2.**
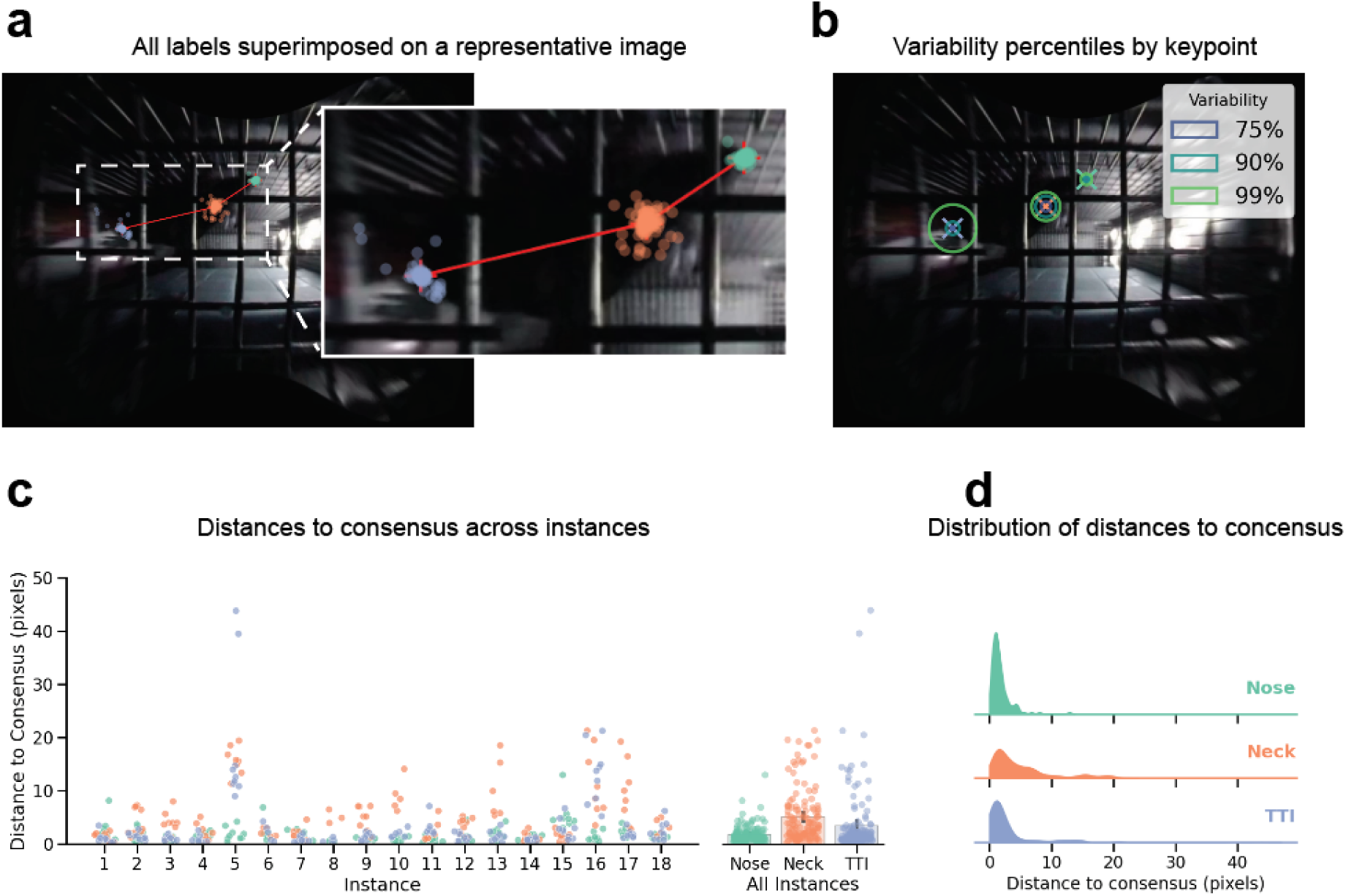
Inter-annotator variability of three keypoint labels. **A.** A representative image with all annotator labels across the full 18 instance set superimposed, corrected for pixel distances. Red crosses represent consensus keypoints. *n* = 144 labels across 18 instances and 15 frames. **B.** Representative image showing localization error as variability percentiles. Consensus labels, represented by crosses, are considered ground truth, and circles represent radial pixel distances of annotator labels in the 75th, 90th and 99th percentiles. **C.** Keypoint labels as distances from consensus in pixels per instance, and across all instances. Bars represent the mean ± the standard error of the mean. **D.** Distributions of error distances to consensus per keypoint.

**Figure 3.**
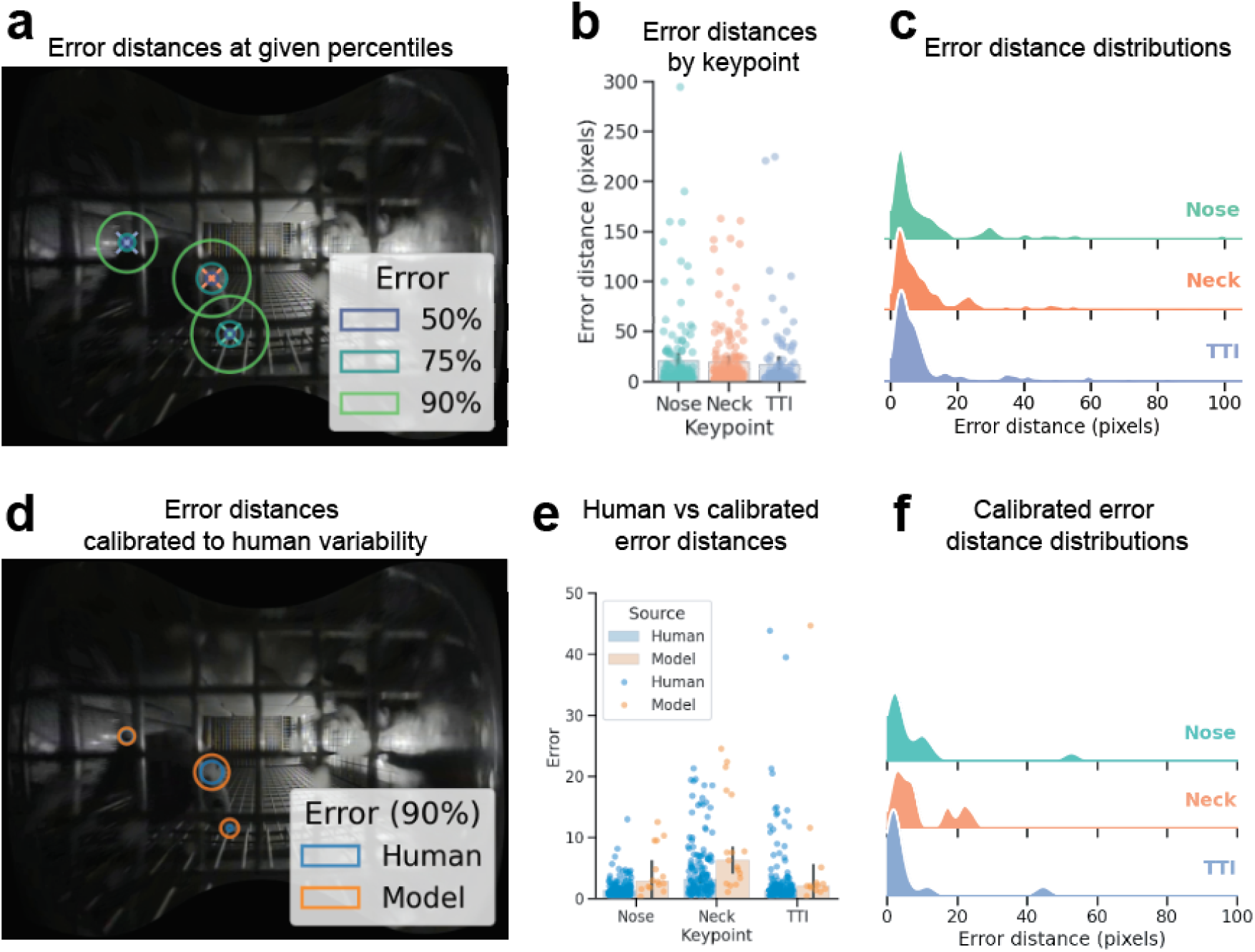
SLEAP pose estimation model performance. **A.** A representative image with radial localization errors overlaid at the 50th, 75th and 90th percentiles, uncalibrated for human variation error. **B.** Uncalibrated localization errors of each keypoint. *n* = 198 held out labeled frames. **C.** Error distances of the SLEAP model in the absence of calibration. **D.** Model localization error and human annotator localization variability at the 90th percentile overlaid as radial distances from consensus in pixels onto a representative image. **E.** Localization error in humans vs the SLEAP model expressed as pixel distances from the consensus. *n* = 144 matched labeled poses. Bars represent the mean ± the standard error of the mean. **F.** Calibrated localization errors of each keypoint. *n* = 198 held out labeled frames.

To provide a scale-invariant measure of accuracy, we computed Object Keypoint Similarity (OKS) scores calibrated to human variability (*32*). Inter-labeler variability yielded scaled standard deviations of 2*σ = 0.2062, 0.2597, and 0.2069, leading to a mean OKS of 0.768. The OKS distribution showed a broad range of multimodal values, with prominent peaks at 0.65–0.7, and 0.8–1 (**Fig. 4a**). Consistent with prior observations of soiling becoming significantly occlusive after two weeks, a longitudinal assessment of average OKS shows distinct decline past day 14 over mission days (**Fig. 4b**).

**Figure 4.**
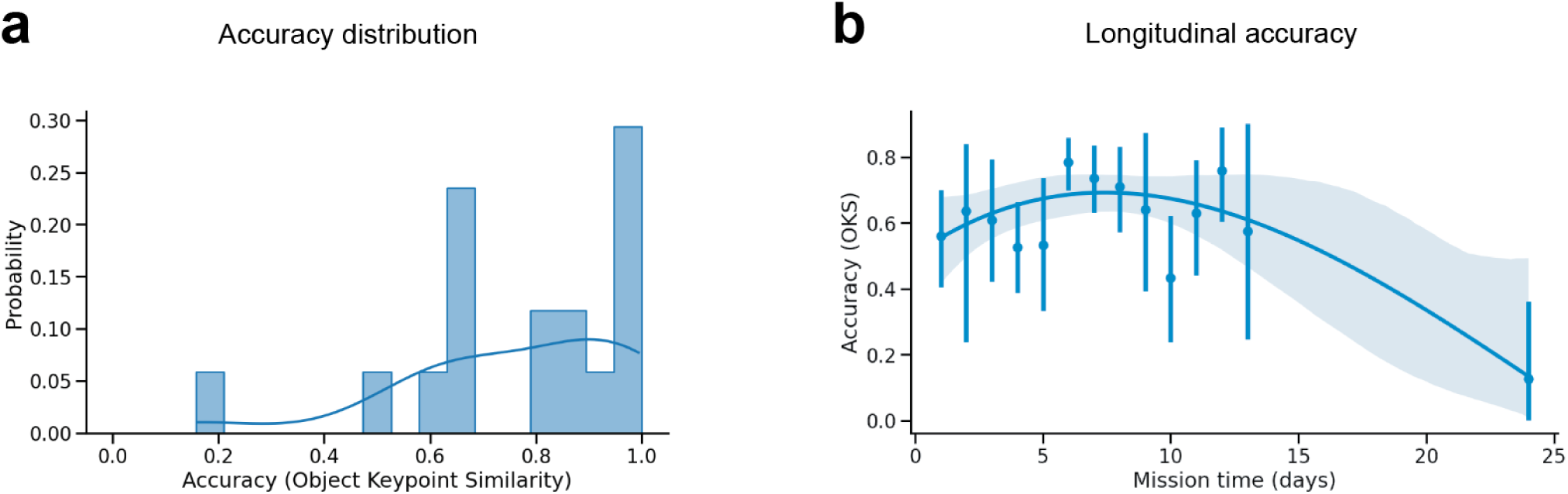
Calibrated SLEAP pose estimation model performance. **A.** A histogram depicting the distribution of Object Keypoint Similarity (OKS) values uncalibrated to human variability. **B.** Average OKS scores across the mission reveals how aggregated soiling occlusions impact predictive accuracy of keypoints. *n* = 3,249 labels. Dots represent the mean ± the standard error of the mean represented by vertical lines. The line represents a regressive fit of average OKS, and the shaded region represents a 95% confidence interval.

### Kinematic Analysis

A distinctive behavioral adaptation that mice display in microgravity is running along the walls of the cage about a central axis. This running behavior is rapid and observed with increased frequency as the mission progresses. Because a major goal of rodent spaceflight research is to understand the mechanistic origin of muscular atrophy and bone demineralization, we sought to understand the centripetal force that mice might be experiencing while circling in a self-centrifugal manner. One key limitation of the monocular camera system of the AEM-X is that there is no overlap of fields of view, and complete loss of some fields of view, making identity tracking of mice infeasible. Therefore, we were limited to performing kinematic analyses only in video segments containing a single subject, which in this dataset was a single mouse on day 10 (**Video S1**). Although only one animal is visible in this clip, mice were group-housed with conspecifics in the AEM-X habitat (see Methods); the remaining animals occupied the adjacent, connected compartment during this recording and were outside the camera’s field of view. Using the centroid of the three keypoints, we first interpolated over frames containing missing predictions. Next, we applied a smoothing filter to account for occasional ‘jumpy’, inaccurate predictions (**Fig. 5a, b**). Once tracking was ethologically accurate, we performed kinematic analysis of circling. We found that velocity oscillates depending on the phase of the circling cycle (**Fig. 5c**). Centripetal acceleration was likewise variable, approximating and occasionally exceeding 1*g* (**Fig. 5d**). To gain a sense of the distribution and dependency of the orbital phase on centripetal force in this sample, we aligned centripetal acceleration estimates by circling phase. Phase-aligned orbital analyses reveal that the majority of circling exerts relatively low centripetal acceleration with values above 1*g* being exceptional (**Fig. 5e**). Because the cage is rectangular, higher forces are exerted on the 90 and 270 degree phases, corresponding to the longer walls (left and right from the camera perspective; **Fig. 5d**).

**Figure 5.**
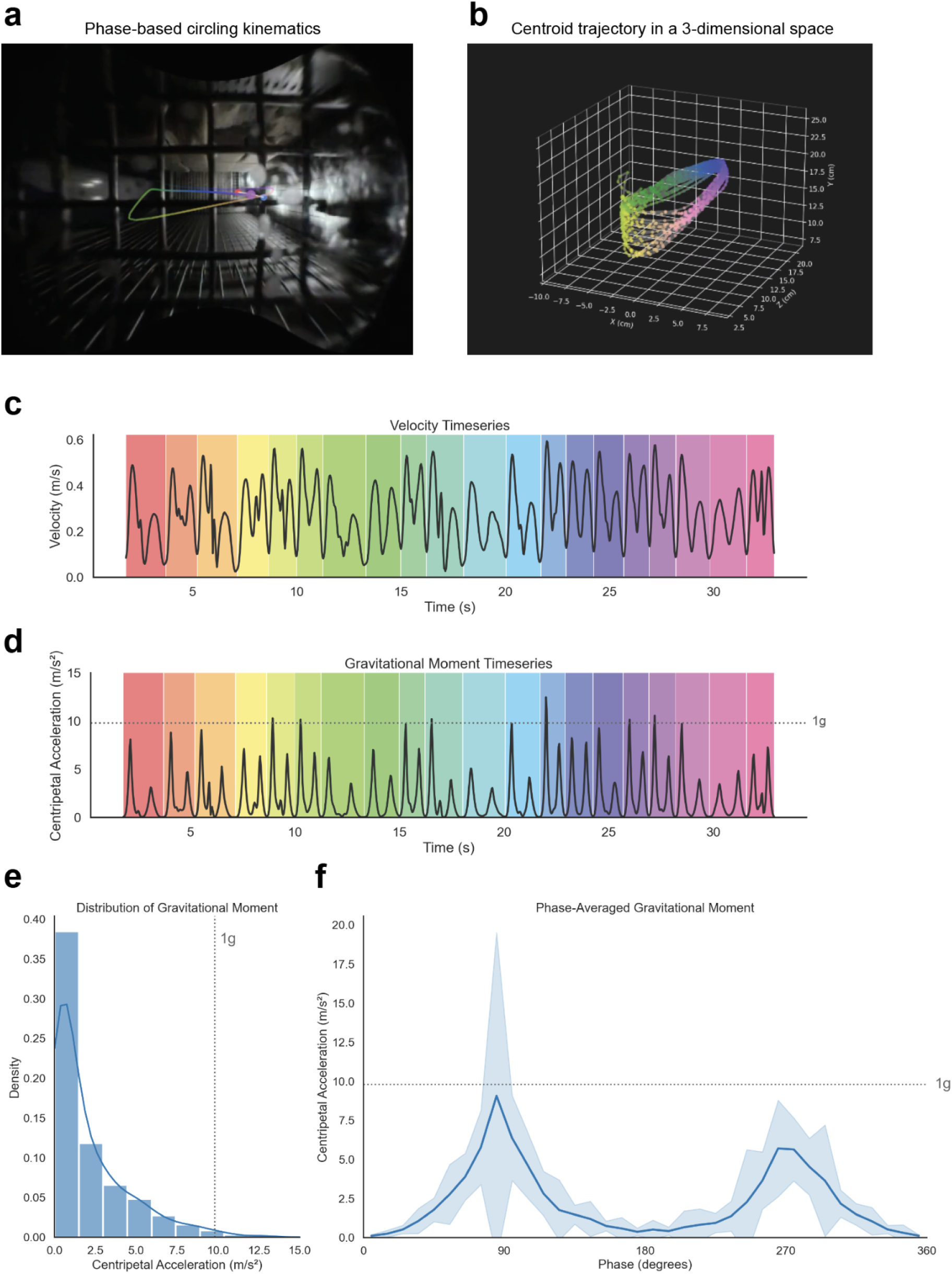
Kinematic analysis of circling behavior of a single mouse on mission day 10. **A.** A still depicting phase-aligned kinematic tracking of circling behavior colormapped to orbit phase. **B.** Reconstructed circling kinematics of a mouse centroid in a 3 dimensional space, colormapped to orbit phase. **C.** Vector- and depth-based velocity estimates during circling in space. **D.** Centripetal acceleration estimates based on phase-aligned orbital tracking and velocity and radial estimates show oscillating centripetal forces that occasionally exceed 1*g*. **E.** Probability of exceeding 1*g* in the video sample by a centripetal force histogram. **F.** Orbit phase-aligned centripetal acceleration estimates show uneven forces throughout the orbit. The line graph represents the mean ± the standard error of the mean. **A-B** colormap represents circling orbit phase, **C-D** colormap represents individual cycles in a select clip of the circling video.

### Behavioral Segmentation

Because pose estimation was limited to three keypoints, a comprehensive skeleton capable of training a behavioral classifier was not possible. Therefore, we sought an alternative behavioral classifier tool, DeepEthogram, which operates at the whole frame level, and uses optic flow to associate motion with a behavioral dataset, and classify behaviors with respect to temporal specificity. The tradeoff with this approach is that, despite having an ability to infer multiple behaviors displayed by multiple mice, this pipeline lacks subject specificity. After studying the dataset, observable behaviors were identified by annotators, and criteria for behavioral annotation were refined for distinction and reproducibility, yielding eight distinct behaviors (including background; **Table S1**). Three trained annotators labeled 411,194 frames across 66 videos spanning the mission, across light cycles and Filter cameras of both cages, yielding 611,045 behavioral event-frames (**Fig. 6a**). Model metrics were collected at the end of the sequencing pipeline. Despite the visual challenges inherent in this dataset, including progressive lens soiling and other occlusions, the model achieved overall accuracy of 0.86 in the training split and 0.90 in the validation split (**Fig. S3a)**. However, performance varied substantially across individual behavioral categories. Mean average precision, a measure of the quality of a model’s ranked predictions via a precision-recall curve, varied across behaviors, ranging from 0.038–0.986 across the test and validation splits (**Fig. S3b-c**). Ethograms comparing manual labels vs predicted behaviors (**Fig. 6a**) depicted a generally overconfident model, owing to its bias towards false positivity.

**Figure 6.**
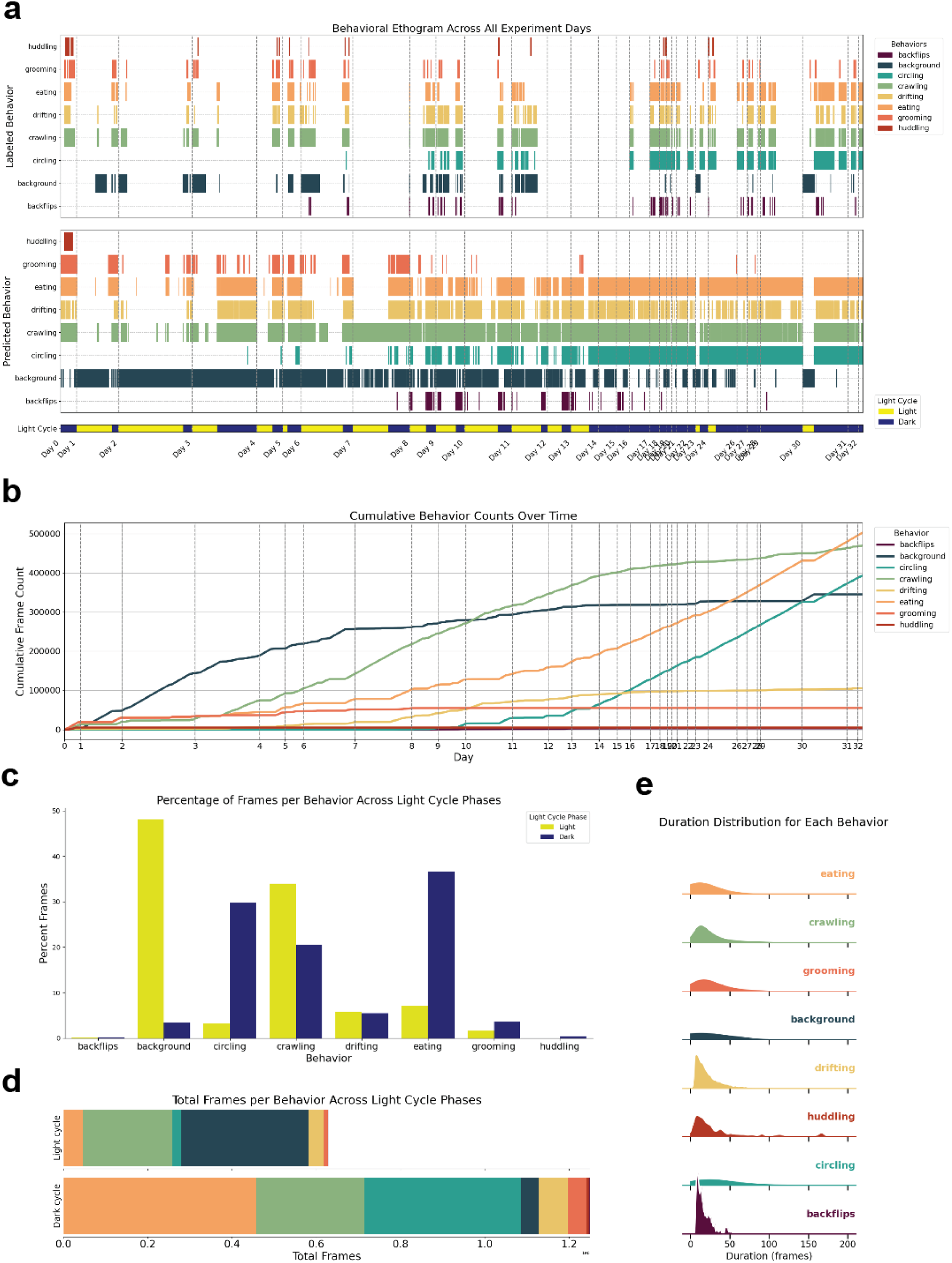
AI-based behavioral segmentation of RR-1. **A.** An ethogram depicting 611,045 human annotated behavior event-frames across 411,194 video frames in 66 videos sampled throughout the mission, used to train DeepEthogram, an optic flow-based behavioral classifier. **B.** A behavioral ethogram of 1,878,357 predicted behavior event-frames across 1,175,485 video frames from 177 videos for the entire RR-1 mission aligned to the light cycle. **C.** Cumulative behavioral inferences by frame across the entire RR-1 mission dataset gives insight into what behaviors might be associated with spaceflight adaptation. Behaviors are color designated, background is a designation for the absence of a detected behavior, including absence of mice in view. **D.** Total inferred frames per behavior across the full mission, aggregated by light cycle. **E.** Distribution of behavioral durations. Most behaviors occur in bouts of <50 frames (∼2s) while circling and background occur over longer, variable durations.

### Behavioral Evaluation

We examined how mission-wide behavioral inferences from our DeepEthogram model might give insight into rodent behavioral adaptations to spaceflight. Because video sampling was uneven across mission days and light cycles, and because the classifier exhibited a false-positive bias, along with highly variable recall metrics, the patterns described below should be interpreted as qualitative trends rather than precise frequency estimates. Model inferences were plotted on a longitudinal ethogram with respect to light cycle (**Fig. 6a**). For better visualization of how behavioral inferences changed with spaceflight duration, cumulative inferences were plotted longitudinally. As noted by others (*25*), mice adapted to crawling along the cage grid as the mission progressed. We qualitatively observed a consistent increase in crawling frame count after day 3 followed by eating, drifting, and circling as the RR-1 mission progressed (**Fig. 6b**). Because video lengths were variable across flight days, inferred behaviors were also expressed as percentage of all inferred behaviors, with respect to light cycle. Circling and eating were behaviors that appeared to be almost exclusively nocturnal. Background, the absence of behavioral inferences, was observed largely during the light cycle, in part due to mice being nocturnally active, but also in part due to many light cycle videos containing no mice in view, as they often sleep in the confines of the Lixit compartment. General locomotion such as crawling and drifting appeared to be even across light cycles (**Fig. 6c**). We next quantified total behaviors inferred in our dataset expressed as total inferred frames. The most commonly observed behavior was eating, followed by crawling, circling, drifting, grooming, huddling, and backflips, though this pattern differed when only considering the light cycle (**Fig. 6d**). To understand this variability in our dataset, we plotted behavioral duration distributions as kernel density plots. Rodent behavior durations are inherently variable and subject to many influencing variables. Crawling, eating, grooming, huddling, drifting, and backflips appeared to occur mostly in the span of 50 frames, or just under 2 seconds. Circling and background behaviors, on the other hand, were far more variable, often lasting 100 or more frames (**Fig. 6e**).

## Discussion

Automated pose estimation and behavioral segmentation of freely behaving mice aboard the ISS proved feasible despite the challenging imaging conditions of the AEM-X vivarium for RR-1. Using SLEAP (*26*), we achieved pose tracking accuracy that approximated human inter-annotator variability across three keypoints, though accuracy degraded with progressive lens soiling after approximately mission day 15. Using DeepEthogram (*33*), we classified eight behavioral categories with high precision and recall overall, although per-behavior performance varied widely and the model exhibited a bias toward false positives that should temper quantitative interpretation of behavioral frequencies. 3D kinematic reconstruction of a prolonged bout of repeated circling was used to estimate centripetal accelerations that periodically approached 1*g*, providing a proof-of-concept for automated biomechanical analysis during spaceflight. Together, these results establish baseline performance metrics for deep learning-based behavioral monitoring in space and identify specific imaging and hardware limitations that constrain current accuracy.

Pose estimation of subjects in complex 3-dimensional poses from a single, fixed monocular camera system is challenging due to inherent self-occlusions of keypoints and varying distances from the camera. Additional occlusions from the internal metal grid, aggregating soiling, and spherical aberration present in the AEM-X vivarium aboard the ISS exacerbate these challenges. We validated the use of deep learning tools against the RR-1 dataset, for which prior manual behavioral scoring and soiling analyses existed (*25*). Some of these challenges can be overcome through computational tools, such as our undistortion protocol (https://github.com/talmolab/spacecage-undistort). In recording systems where subjects have sufficient trackable keypoints, tools exist to convert monocular tracking into 3-dimensional poses (*34–39*); however, keypoint density was insufficient in this dataset for such approaches. We explored a tool for removing the metal grid from the video (*40*) but found that outputs yielded artifacts, lowered overall resolution, and did not sufficiently recover animal body parts behind the grid, making the approach incompatible with our pose estimation and behavioral segmentation pipelines. Subsequent iterations of grid-removal methods may overcome this limitation. Prior work tracked mouse activity in the JAXA Habitat Cage Unit using conventional, non-deep-learning tracking tools (*28*), and pose estimation has recently been applied to *C. elegans*, *Drosophila*, and zebrafish on the Chinese Space Station, Tiangong (*27*). The present study is, to our knowledge, the first to apply deep learning-based pose estimation and behavioral segmentation to establish the feasibility of computer vision-based phenotyping of rodent behavior in spaceflight.

One of the goals of pose estimation in our study was to build a kinematics-based behavioral classifier. However, the video quality, numerous occlusions, and absence of depth-sensing and overlapping cameras meant that very few body points could be reliably and reproducibly labeled from all 3-dimensional poses of a mouse as captured by a monocular camera system. Therefore, we relied on a behavioral classifier pipeline that instead uses optic flow features at the cost of loss of individual tracking and true attribution to kinematics. Using this pipeline, behavioral inferences across the RR-1 mission recapitulated and extended findings from prior manual scoring by Ronca et al. (*25*), who analyzed video from both the RR-1 NASA Space Biology Validation study (the same video as this study), and a parallel ISS National Lab mission in the same RR-1 payload. The present study analyzed only NASA Space Biology RR-1 Validation footage, so direct comparisons to their behavioral frequencies are limited. The conclusions that can be drawn from our behavioral analyses are also limited due to severe sampling biases of video duration and light cycle, where light cycle videos are captured at a greater length, but almost exclusively until mission day 13. Model-based behavioral precision and recall are also specific to individual behaviors, with huddling and backflips having distinctly poor metrics in part due to huddling labels involving very little movement, yielding low optic flow features, and in part due to the very few backflip and huddling examples available in the labeled data. Despite these limitations, data output by this model offers insight into rodent behavioral adaptations to spaceflight. Locomotor behaviors stand out as showing the most temporally distinct changes. We observe a drastic increase in crawling frequency after the first several mission days, consistent with the previously reported shift from forelimb to quadrupedal locomotion as mice adapted to microgravity. Eating detections were the next distinct rise, although this appears to be more aligned to light cycle sampling availability, as eating is observed almost exclusively in the dark cycle, when mice, which are nocturnal prey species, are more active. We next observed increased drifting, which we speculate reflects mice learning a more efficient way to move, or perhaps a play behavior. Circling detections increased toward the second half of the mission, and backflips emerged as an increasingly frequent behavior over time, both of which also aligned with the dark cycle and can be considered play behaviors. Early in the mission, drifting was commonly observed as mice collided in different trajectories; the progressive decline in drifting detections suggests that mice learned to navigate the microgravity environment by maintaining contact with cage surfaces. Background detections, which reflect the absence of visible mice in the camera field rather than a scored behavior, occurred predominantly during the light cycle, while circling and eating were observed almost exclusively during the dark cycle, consistent with expected nocturnal activity patterns in mice. However, these circadian patterns must be interpreted cautiously: light-cycle videos were archived almost exclusively before mission day 13 due to thermal issues that led to discontinuation of light-cycle captures (*7*), so apparent light/dark differences are confounded with mission phase and cannot be attributed to circadian rhythmicity alone. This analysis also lacks ground control behavioral comparisons; behavioral trends described here as spaceflight adaptation could partly reflect habituation to a new housing environment independent of microgravity. All behavioral frequencies reported here derive from model inferences with a known false-positive bias, and per-behavior classifier performance varied widely (mAP range: 0.04-0.99); these results should therefore be treated as qualitative trends rather than precise quantitative estimates.

Although pose estimation was limited to three reliable keypoints, kinematic analyses are still possible in some use cases. Due to the common focus of flight experiments on muscular atrophy and skeletal remodeling, we sought to estimate the mechanical loading forces mice experience during circling. Individual pose-based tracking proved extremely difficult in a monocular camera system with subject-based occlusions. Therefore, kinematic reconstruction of circling behavior depended on the only sample available with just one subject in view. This sample was captured on mission day 10 (15 days from launch), and we estimated centripetal accelerations that periodically approached and slightly exceeded 1*g*. These estimates derive from a single circling bout of one mouse on one mission day and should be interpreted as a proof-of-concept demonstration rather than a generalizable measurement. Circling speed likely increases as the mission progresses based on qualitative observation, suggesting that later-mission circling bouts may produce higher centripetal forces, though we lack additional kinematic data to confirm this. Whether self-generated centripetal loading during circling provides a meaningful countermeasure to microgravity-induced musculoskeletal deconditioning remains an open question. Spaceflight-induced muscular atrophy was documented in mice from the same RR-1 mission counterpart by the ISS National Lab portion (for which videos are not available), where even MuRF1 knockout mice were not protected from muscle loss (*41*). Mice returned from RR-9 in 2017 showed altered gait characteristics and spaceflight-induced articular cartilage loss in the tibial-femoral joint (*42*, *43*), suggesting that the mechanical loading mice experience in the AEM-X vivarium is insufficient to prevent musculoskeletal deterioration. By contrast, mice housed in 2023 in JAXA’s Mouse Habitat Unit-8 centrifugal vivarium and subjected to 1*g* for 32 days did not show maladaptive gait upon return (*44*), a result that supports artificial gravity as a potential countermeasure for musculoskeletal deconditioning during Moon and Mars missions. The circling of freely behaving mice in the AEM-X vivarium appears insufficient as a countermeasure to spaceflight-induced gait alterations. Future iterations of the vivarium system should include multiple cameras with overlapping field of view to overcome self- and subject occlusions, leading to more reliable tracking and kinematic analyses.

As the first NASA rodent mission to the ISS, RR-1 served as a validation mission for the AEM-X vivarium system (*7*) and generated a broad body of downstream science through NASA’s open science infrastructure. The Biospecimen Sharing Program and the NASA Biological Institutional Scientific Collection enabled tissue sharing-based studies spanning multiple rodent organ systems (*17*, *45–52*) while data sharing, data standardization, and analysis-visualization tool development efforts through the NASA Open Science Data Repository(*30*), Ames Life Sciences Data Archive, and GeneLab (*53*) supported numerous omics and phenotypic-physiological analyses from this same mission (*54–63*). Among these, one study found that brains from the mice in the present study showed lower expression of inflammatory cytokines relative to ground controls (*17*), raising the question of whether neuroinflammatory changes relate to the emergence of circling behavior, though the present study cannot establish such a link. Integrating the video-derived behavioral phenotypes reported here with the rich omics and physiological datasets already available from RR-1 through OSDR could enable multi-modal approaches such as anomaly detection, temporal pattern mining, and predictive modeling to identify molecular correlates of behavioral adaptation to spaceflight.

Several limitations constrain the conclusions that can be drawn from this work. The RR-1 video archive was designed for animal health monitoring, not behavioral research. Because RR-1 was in part a validation experiment, unforeseen hardware issues resulted in inconsistent sampling across mission days, light cycles, cages, and camera views (*7*). Light cycle video capture was discontinued after mission day 13, preventing analysis of circadian behavioral patterns across the full mission. Progressive lens soiling degraded pose estimation accuracy after approximately day 15, and the internal metal grid occluded keypoints throughout the mission (**Table 1**). The behavioral classifier showed mixed performance, which was highly dependent on behavior, and therefore downstream behavioral frequency analyses should be treated as qualitative. The kinematic analysis of circling was limited to a single mouse in a single video on mission day 10. We did not analyze ground control videos in this study, precluding direct flight-versus-ground behavioral comparisons. Individual mice could not be distinguished due to the absence of visible identifying markers, preventing individual-level behavioral tracking, as also noted by Ronca et al. (*25*). Finally, due to the brief duration of this mission, mice absorbed an estimated 8.6–13 mGy of whole-body radiation during the 37-day mission as mined from OSDR’s RadLab (**Fig. S1**) (*64*), well below the average ISS astronaut dose of 28 mGy (*65*) and far below anticipated doses for lunar or Mars missions (*66*). The contribution of this low dose to the behavioral patterns observed, if any, cannot be determined from this study.

**Table 1.**
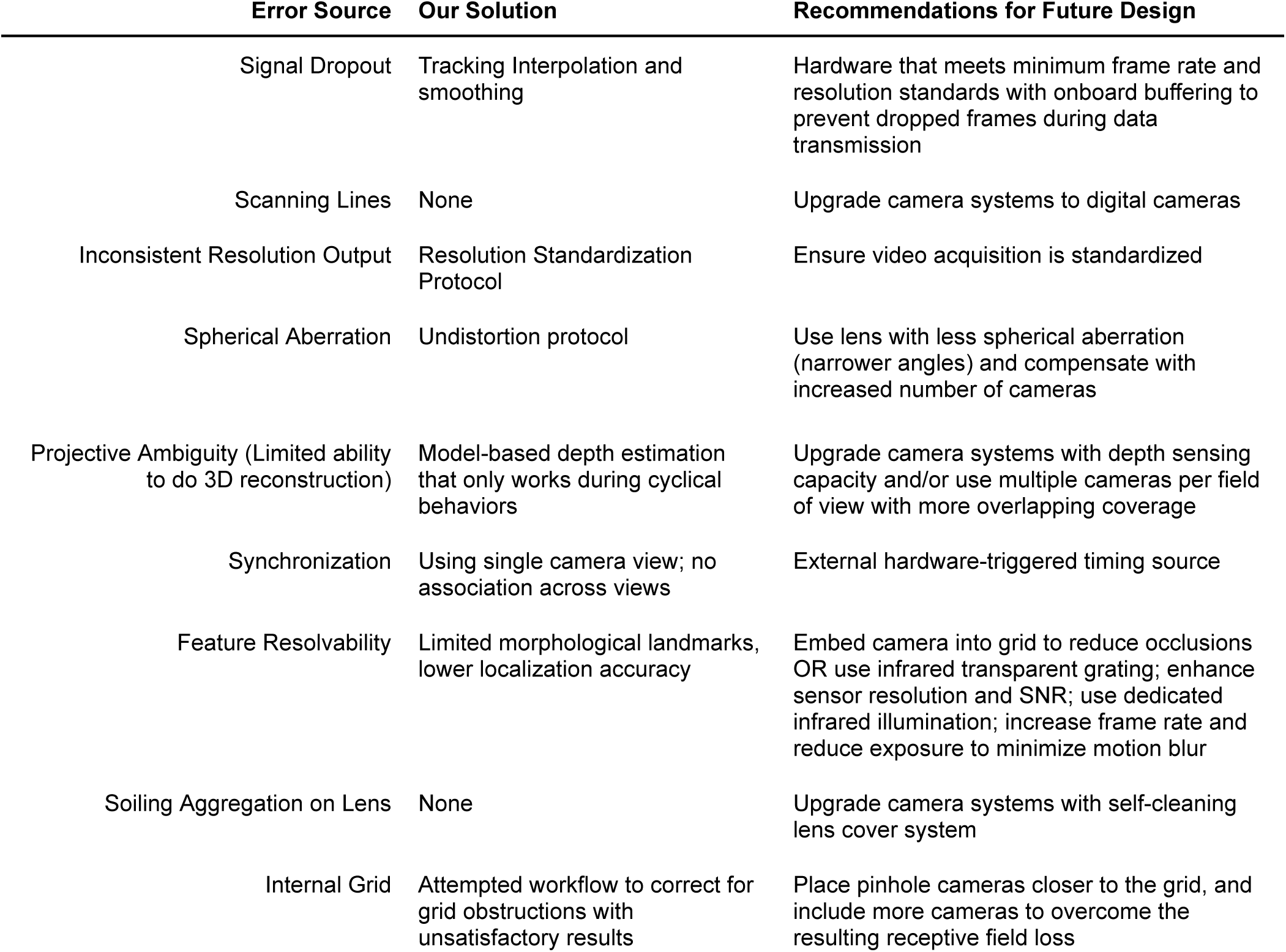
Obstacles to video-based motion capture in the AEM-X vivarium, with workflow solutions and recommendations for future hardware iterations. Each row describes an imaging or acquisition challenge encountered in the archival RR-1 video dataset, the mitigation applied in the present analysis, and a hardware or protocol recommendation to overcome the challenge in future Rodent Research missions or research vivariums.

One advantage of the AEM-X vivarium as a repeatable experimental arena with a static background is that the pose and behavioral models from this work can be applied to archival video from other Rodent Research missions with minimal additional labeling. One caveat is the introduction of environmental enrichment huts beginning with RR-5, which will require an additional labeled dataset to account for the changed background and potential occlusion of circling behavior, though fine-tuning on top of the models presented here should require far fewer labels than training from scratch. The models may also serve as starting points for pose and behavioral segmentation in other spaceflight vivaria, including the Bion M-1, the Mouse Drawer System, the Space Shuttle AEM, and the JAXA Mouse Habitat Cage Unit (*8*). It also could inform the design of behavioral monitoring systems for future vivaria aboard lunar surface habitats or Mars transit vehicles. Automated behavioral analysis reduces the burden of manual video scoring by mission support staff, a consideration that becomes increasingly important as missions extend beyond low Earth orbit where real-time ground support is constrained by communication delays. More broadly, this work demonstrates that AI-based behavioral segmentation and kinematic extraction are feasible with archived spaceflight video, opening the possibility of retrospective behavioral analyses across the more than 70 rodent missions that have flown to space (*28*). Such capabilities align with broader efforts to integrate AI-based biomonitoring and maximally autonomous research platforms into deep space mission operations (*67*, *68*).

The need for automated longitudinal behavioral monitoring extends beyond animal models. Behavioral monitoring can provide an abundance of performance information such as retroactive analysis of work efficiency, and interpersonal activity, yielding behavioral biomarkers indicative of operational or psychiatric risk (*29*, *69*). Potential mission critical risk mitigation tools and other countermeasures such as fatigue assessment, psychiatric condition alert, and mood evaluation could arise from the use of AI (*67*, *68*). To our knowledge, no published work has applied deep learning-based video pose estimation or automated behavioral segmentation to human crew in spaceflight or analog environments; current crew behavioral monitoring relies on self-report questionnaires, wrist actigraphy (*70*), and periodic neurobehavioral testing (*71*). Because crew social dynamics, team function, and conflict mitigation are areas of key research focus, AI-based behavioral tools can improve social dynamic measures in analog and space environments (*20*, *72*, *73*). Rodent behavioral models that capture longitudinal changes in activity, locomotion, and social dynamics under spaceflight conditions represent a translational step toward analogous crew monitoring systems.

The labeled datasets, trained models, and analysis code are publicly available through NASA OSDR at OSD-952 and through GitHub, enabling the spaceflight biology community to build on and improve these methods. This work establishes a proof of concept that AI-based behavioral segmentation and pose-driven kinematics can be extracted from archived spaceflight video originally collected for other purposes. The models and labeled datasets produced here are ready to be applied to or improved by additional Rodent Research missions, and the analytical framework can be extended to other model organisms and vivarium systems as spaceflight biology expands beyond low-Earth orbit. We also identify challenges to AI-based behavioral phenotyping and kinematic analyses in current hardware, propose provisional solutions to some of these obstacles, and recommend modifications needed in future flight hardware iterations (**Table 1**). As NASA and both its commercial and international partners prepare for exploration-class missions to the Moon and beyond, automated behavioral phenotyping systems that overcome the imaging limitations identified here will lay the groundwork for continuous, Earth-independent monitoring of animal and, ultimately, crew behavioral health during long-duration spaceflight.

## Materials and Methods

### Mice

All mouse housing conditions, flight, and hardware information were previously reported (*7*, *25*). In brief, all subjects in the videos were female C57BL/6J (The Jackson Laboratory, Bar Harbor, ME), group-housed with conspecifics in the AEM-X habitat, which consists of two interconnected compartments (*n* = 5 mice per cage). Mice were 16 weeks of age at the time of launch. Mice received deionized, autoclaved water, and NASA Type 12 Nutrient-Upgraded Rodent Food Bars *ad libitum*. During launch and transit to the ISS, mice were housed in Transporters as described previously (*7*, *25*); all video analyzed in the present study was collected after transfer to the AEM-X habitat aboard the ISS. A parallel ground control group was concurrently housed on the ground in identical flight hardware (*7*, *25*) but was not included in the present video analyses. Mice were kept on a 12h/12h light/dark cycle (0600–1800 GMT light) (*74*). All procedures and experiments were approved by the NASA IACUC and International Space Station National Laboratory IACUC, and conformed to relevant federal regulations of research animal use (NRC Guide for the Care and Use of Laboratory Animals and Title XIV) as previously described (*7*, *25*). Mice experienced approximately 8.6-13 mGy of whole-body space radiation for the duration of the mission (**Fig. S1**). All videos and associated flight data were acquired through a data request from NASA’s OSDR and ALSDA.

### Flight and Hardware

Mice in this study were housed in the AEM-X vivarium and transport system as previously described (*7*). All footage was captured in the AEM-X vivarium. RR-1 launched on September 21, 2014 from Kennedy Space Center (SpaceX-4), and arrived at the ISS four days later, at which point cages were transferred to their enclosure module racks. The rodent habitats contained four cameras pointing to two different views of the Lixits (Pinhole WD Color Model PC315XP), and two views of the filter area (B/W Micro Video Model PC206XP, Austin, TX). Data were digitally routed from the ISS through the Huntsville Operations Support Center at Marshall Space Flight Center, to the Multi-Mission Operations Center server at Ames Research Center for rodent health check analysis by RR-1 team staff. Videos were then securely stored by ALSDA (part of OSDR) on local servers, although later RR missions would refine this workflow to a near real-time and remote solution including a secure Collaborative Life Sciences Repository enabling veterinarians, engineers, Principal Investigators, and RR science teams to check on animal health and hardware functioning in accordance with IACUC and mission requirements (IACUC #NAS-13-002-Y1). Video captures were originally intended for health evaluations, and programmed to upload one-hour videos at the start of the light cycle and at the start of the dark cycle daily. However, thermal issues during recording necessitated variations in video capture schedules, lengths, and resolutions, eventually leading to discontinuation of light cycle captures (*7*). Video capture began on flight day 5 and lasted through flight day 37, and were captured between 15 minutes and 2 hours following the start of a light cycle phase. Timestamps and recording dates conform to the ISS National Laboratory’s GMT+0 timezone. The first video capture day (day 0) reported in this experiment corresponds to flight day 5.

### Video Preprocessing

The RR-1 video files were concatenated camera views of highly variable scheduled captures, where camera views cycled through the four cameras of each side of the AEM-X rodent vivarium aboard the ISS. Behavioral and keypoint scoring was extremely challenging with Lixit cameras due to the proximity to animals. Therefore, like prior work, only the filter cameras were used for analyses (*n* = 354). To isolate individual camera views, and to omit Lixit views, videos were manually cropped at the change of each camera field. Of the 354 archived flight and ground filter videos, only 177 were flight videos of suitable quality. Mann-Whitney tests were employed to compare the distributions of total frames and sampling days among Dark and Light Cycle Flight Filter videos. Distribution comparisons of sampling days and frames per sample were visualized as Gardner-Altman plots (**Fig. S2c**).

### Video Normalization

The RR-1 archive presented several video quality challenges. To minimize bias in computer vision-based approaches to pose estimation and behavioral segmentation, we employed two normalization strategies. Most flight filter videos had a resolution of 640 x 480 pixels or higher, with a few exceptions, but all at 30 frames per second (**Fig. S2a**). Therefore, we first wrote a custom script to center and standardize all videos to 640 x 480 pixels, effectively maintaining even pixel size, aspect ratio, and field of view across all video samples by downsampling or padding videos based on a common middle point (**Fig. 1a**). Next, we created a pipeline for correcting for the spherical aberration of the camera lenses by using the 1 cm x 1 cm metal grids in the field of view as calibration references to estimate distortion coefficients (https://github.com/talmolab/spacecage-undistort).

### Human Keypoint Labeling Consistency Validation

To assess how labeling might be affected by human variation, nine labelers were trained to label the snout, neck, head, left ear, right ear, left shoulder, right shoulder, left haunch, right haunch, trunk, TTI, the tail tip, and three equidistant points along the tail between the TTI and tip along the anatomical localizations of the mouse skeleton. The complex poses and high variability in angles quickly ruled out all body points with the exception of the snout, neck, and TTI. Labelers used the SLEAP graphical user interface on a frame-by-frame basis. To ensure labeling reproducibility, we conducted variability analyses that compared the same three keypoints across 18 frames annotated by all nine labelers, yielding 144 labels per human annotator. The selected frames varied widely in animal pose and proximity to the camera, and across nine spatial quadrants of the frame. To establish a human baseline for keypoint localization accuracy, we first established a consensus keypoint as the pixel coordinate average of all human labels per keypoint. Next, variability was calculated as pixel distances from the consensus point, and expressed as the OKS (*32*). Inter-labeler variability was visualized at the 75th, 90th and 99th percentiles.

### Pose Estimation Model Training and Evaluation

Nine human annotators labeled a representative dataset containing videos across the mission to account for lens occlusions due to soiling. *n* = 3,249 labels span *n* = 2,063 frames of *n* = 78 videos throughout the mission. The labeled keypoints included the snout, neck and TTI. The SLEAP model was trained with a bottom-up backbone, in grayscale. Inferences were performed on *n* = 177 videos. SLEAP performs pose estimation independently on each frame without temporal context; temporal information was used only in the downstream kinematic reconstruction of circling behavior. The complete label dataset, model, training parameters and inferences are provided in the OSDR repository OSD-952. Model evaluation was first performed on a held-out test set to assess localization error per keypoint, then calibrated against human inter-annotator variability using OKS scores (*32*).

### Behavioral Segmentation Model Training and Evaluation

Due to visual occlusion and a dynamic 3D environment, we chose the open-source Python-based package DeepEthogram as the predictive model. DeepEthogram is a pipeline of three separate models that work in concert to predict behavior. First, it uses a MotionNet-based flow generator to assess optic flow-based movement-related features. Next, it trains the feature extractor, a large convolutional neural network that associates individual labeled behaviors within individual timepoints by compressing snippets of optic flow and individual still images into a lower dimensional set of features. This classifier effectively learns the features of an animal’s body that are relevant to behaviors. Lastly, DeepEthogram trains a sequencing classifier that takes relevant, compressed behavioral features across many frames to learn and infer the onset and end of behaviors (*33*). We first established behavioral criteria based on the literature, and refined them to ensure reproducibility among behaviorists. Behavioral categories and their operational definitions are provided in **Table S1**. Three trained behaviorists labeled a total of *n* = 411,194 frames across *n* = 66 videos throughout the entire experiment duration, across both cages and the light and dark cycles. Behavioral labeling was done on DeepEthogram’s graphical user interface on a frame-by-frame basis (**Fig. 1d**). DeepEthogram does not rely on human-annotated poses to infer behaviors. Therefore, multiple behaviors can be labeled and inferred within any given frame. To optimize training accuracy, we used DeepEthogram’s slowest and most accurate pre-weighted models, TinyMotionNet3D for the flow generator, and 3D-ResNet34 for the feature extractor/sequence. Model performance was evaluated with a test split of *n* = 335,902 frames and a validation split of *n* = 75,292. All processed flight filter camera videos were inferred (*n* = 177 videos; total frames = 1,175,485).

### Circling Analysis

Circling is a behavioral adaptation of mice to microgravity previously described (*25*). This behavior involves mice running on four perpendicular walls about a central axis. Because one of the goals of rodent spaceflight research is to examine microgravity-induced bone demineralization and muscular atrophy, it is critical to understand the musculoskeletal load that rodents experience in the RR missions. To this end we devised an analysis pipeline that factors the dimensions of the AEM-X vivarium (*75*), and used pose inferences from our SLEAP model on the only video sample of a single mouse circling, with tracklet interpolation (three-frame median filter) and smoothing (Gaussian; σ=1.5) procedure. We then estimated z-plane depth based on body size, using the 5th–95th percentile of the body length to omit outliers and scale pixel sizes to actual distances. We then reconstructed the trajectory of the mouse applying wall constraints based on cage dimensions, such that keypoints stay within 2 cm of the 3-dimensional construct of the walls. Next, to determine the directionality of circling based on movement orientation, we drew a centerline along the video, marking proximal and distal crossings. Knowing the orientation of circling, we segmented the laps based on the trajectory of crossings. With these 3-dimensional kinematics, we generated a time series of velocity, acceleration, and radius of curvature. We applied these kinematics to estimate centripetal acceleration:

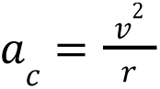

### Radiation and Other Physico-Chemical Environmental Telemetry

Whole-body radiation dose estimates for the RR-1 mission were derived from two ISS dosimeters queried through NASA OSDR’s RadLab portal (https://visualization.osdr.nasa.gov/radlab/)(64). The Radiation Environment Monitor (REM; instrument ID: REM-LAB103-pre) and the Tissue Equivalent Proportional Counter (TEPC; instrument ID: TEPC-SMP327-pre) provided absorbed dose rate measurements spanning the mission duration from September 21 to November 2, 2014 (**Fig. S1a**). Mean absorbed dose rates were approximately 8.5 μGy/hour (REM) and 12.9 μGy/hour (TEPC). Cumulative absorbed doses were estimated at approximately 8.6 mGy (REM) and 13.0 mGy (TEPC) over the 37-day mission. The discrepancy between instruments reflects differences in detector sensitivity, shielding geometry at placement, and dose equivalence vs raw dose estimates; radiation dose estimates to biological payloads carry approximately ±10% uncertainty because most flight experiments do not employ dedicated integrated dosimeters adjacent to the vivarium. ISS Columbus module environmental telemetry data including temperature, relative humidity and CO_2_ concentration for the ISS flight hardware were obtained from NASA OSDR’s Environmental Data Application (https://visualization.osdr.nasa.gov/eda/) (*30*, *31*) and are shown in **Fig. S1b–d.**

## Acknowledgements

This work was funded by NASA Grant Number 80NSSC24K0346 through the NASA Human Research Program (HRP, PI: TDP) and NIH Grant Number R01MH129970 (NIMH, PI: AJE). The authors thank the NASA Open Science Data Repository and Ames Life Sciences Data Archive teams for supporting the acquisition of the RR-1 video data, answering questions about the RR-1 payload, and curating the results and outputs from this study into OSD-952. In particular, we thank Rachel Gilbert, Danielle Lopez, Jonathan Galazka, and Samrawit Gebre. We also thank the NASA Biological and Physical Sciences Division for supporting OSDR and ALSDA which are valuable national and global resources for the future of space biology and space health. We also thank HRP’s Human Factors and Behavioral Performance element for supporting this work over the course of the grant. Thanks also to Ryan Fisher and Vandana Verma for their communications on the RR AEM hardware. RTS is part of the “AI for Life in Space” (AI4LS) group of researchers at NASA Ames, and grateful for the support from them as well as the NASA Science Mission Directorate’s ‘Foundation Model’ effort, specifically for the NASA Biological and Physical Sciences division. Anthropic’s Claude was used to assist with editing during manuscript preparation. The authors take full responsibility for all content and verified everything for accuracy.

## Author Contributions

F.C.K., T.D.P., A.J.E., L.M.S. and R.T.S. conceptualized the study. T.D.P., F.C.K., M.T.M., A.M.III, M.M., S.G., J.H., M.B., S.J. and A. Mahajan curated the data. T.D.P., F.C.K. and M.T.M. performed formal analysis. T.D.P., L.M.S., R.T.S. and A.J.E. acquired funding. A.J.E., F.C.K. and T.D.P. administered the project. F.C.K. and T.D.P. developed the software and code. F.C.K. and T.D.P. created the visualizations. A.J.E., F.C.K., T.D.P., L.M.S. and R.T.S. wrote the original draft. A.J.E., F.C.K., T.D.P. and R.T.S. reviewed and edited the manuscript. These authors jointly supervised this work: F.C.K., T.D.P. and A.J.E.

## Data Availability

All transformed videos, annotated datasets, model files, and inferences are available on NASA’s OSDR at OSD-952 (https://doi.org/10.26030/6m3b-gd71); files containing embedded rodent imagery-video will require approval from the NASA OSDR team prior to access, while files without embedded rodent imagery are freely available. The Radiation and Environmental Telemetry can be found in OSDR’s RadLab portal (https://visualization.osdr.nasa.gov/radlab/) and OSDR’s Environmental Data Application (https://visualization.osdr.nasa.gov/eda/).

## Code Availability

Analysis scripts and workflows are available at https://github.com/talmolab/Space-SLEAP, and the video transformation and spherical aberration correction pipeline is available at https://github.com/talmolab/spacecage-undistort. Links to these GitHub repositories are also linked at OSD-952.

## Supplementary Figures

**Figure S1.**
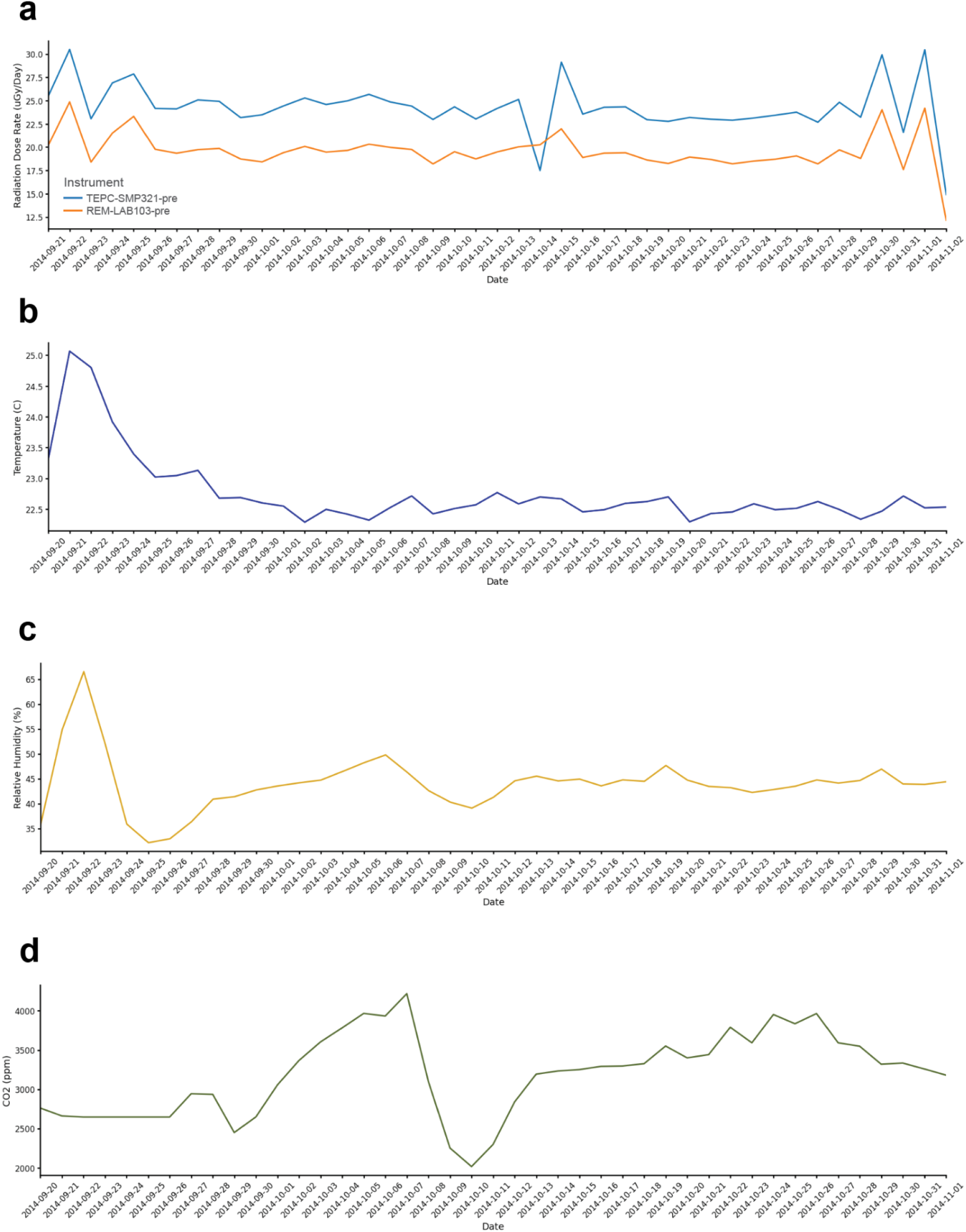
RR-1 telemetry for **A.** radiation dosimeters, **B.** temperature, **C.** relative humidity, and **D.** CO_2_ concentration in the Columbus module of the ISS.

**Figure S2.**
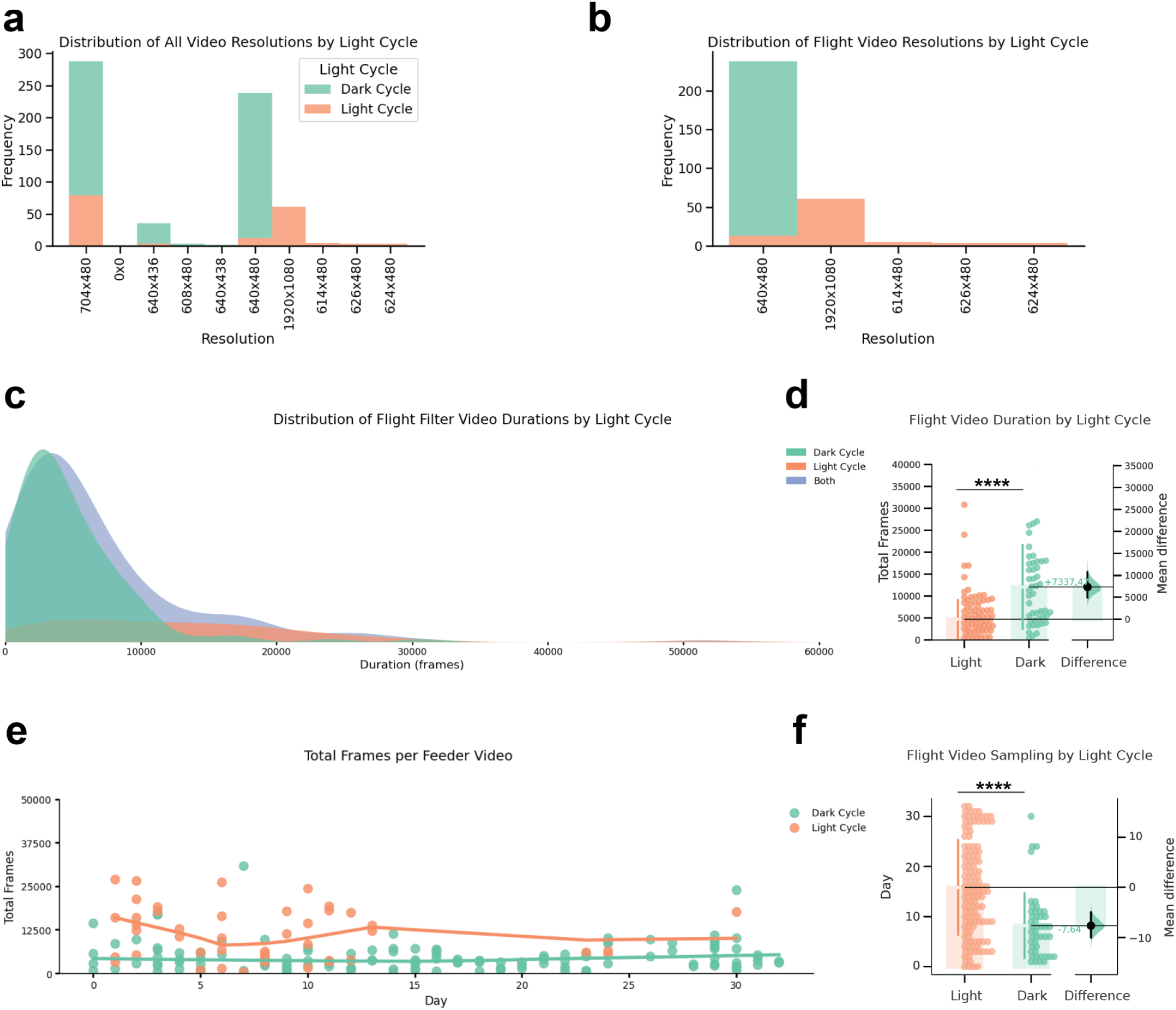
RR-1 Filter video dataset distributions. **A.** Distribution of resolutions of unprocessed RR-1 Ground and Flight video dataset. **B.** Distribution of resolutions of unprocessed RR-1 Flight videos. **C.** Distribution of Flight video durations. **D.** Comparison of Flight video durations by light cycle. **E.** Distribution of Flight video duration by day and light cycle. **F.** Comparison of Flight video day sampling by light cycle. Bars represent the mean ± the standard error of the mean; Day and frame sampling comparisons performed with the Mann-Whitney test; **** *P* < 0.0001

**Figure S3.**
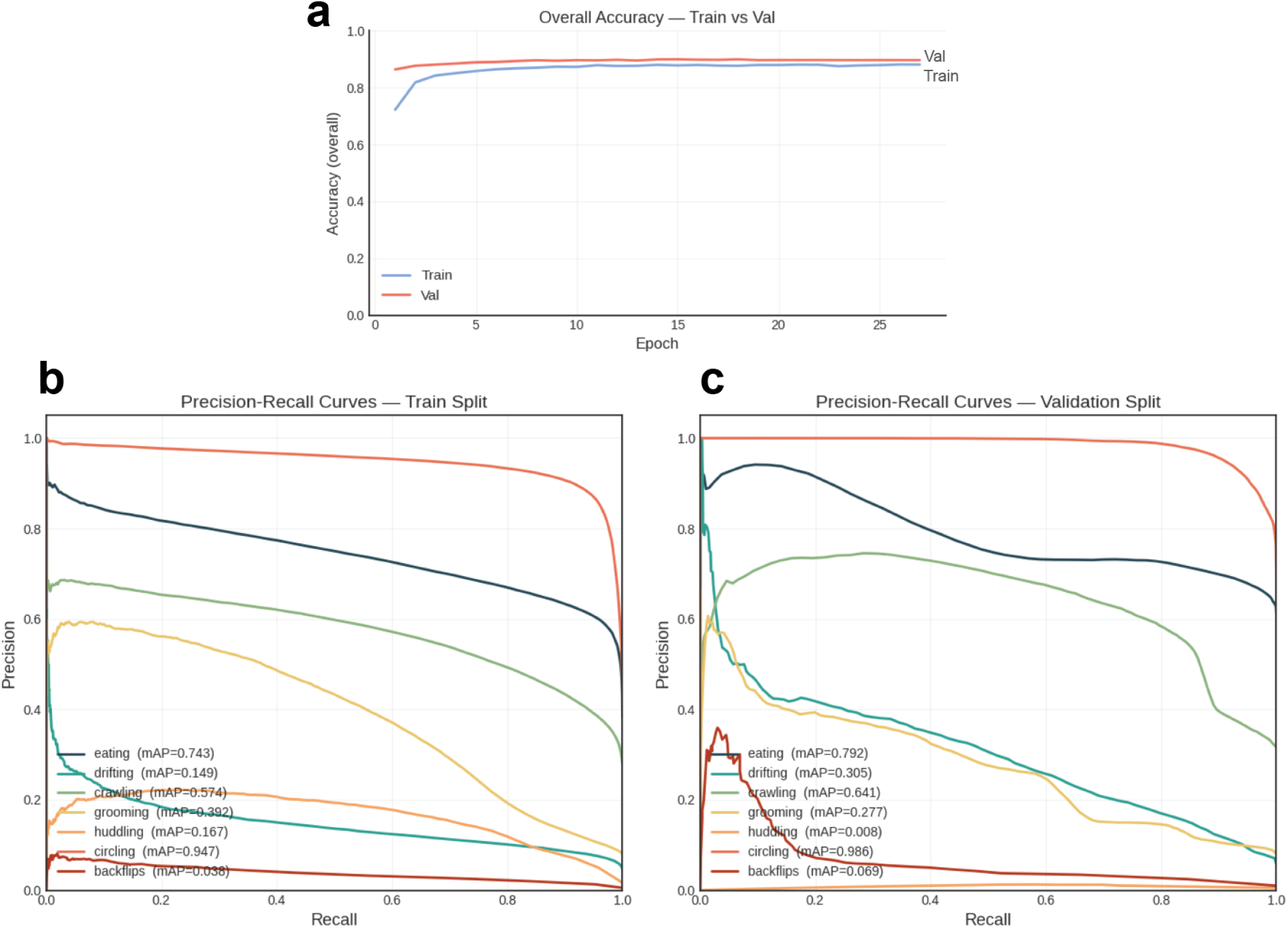
DeepEthogram model performance. **A.** Accuracy for all behaviors in the Validation and Train splits at the end of the Sequence model. **B.** Precision-Recall curves and mean average precision values (mAP) for each behavior in the Train split. **C.** Precision-Recall curves and mAP values for each behavior in the Validation split.

**Supplementary Movie 1.** Animation of inferred 3-dimensional kinematic trajectories, and real-time phase-based centripetal acceleration estimation of centroid during a bout of repeated circling behavior.

**Table S1.**
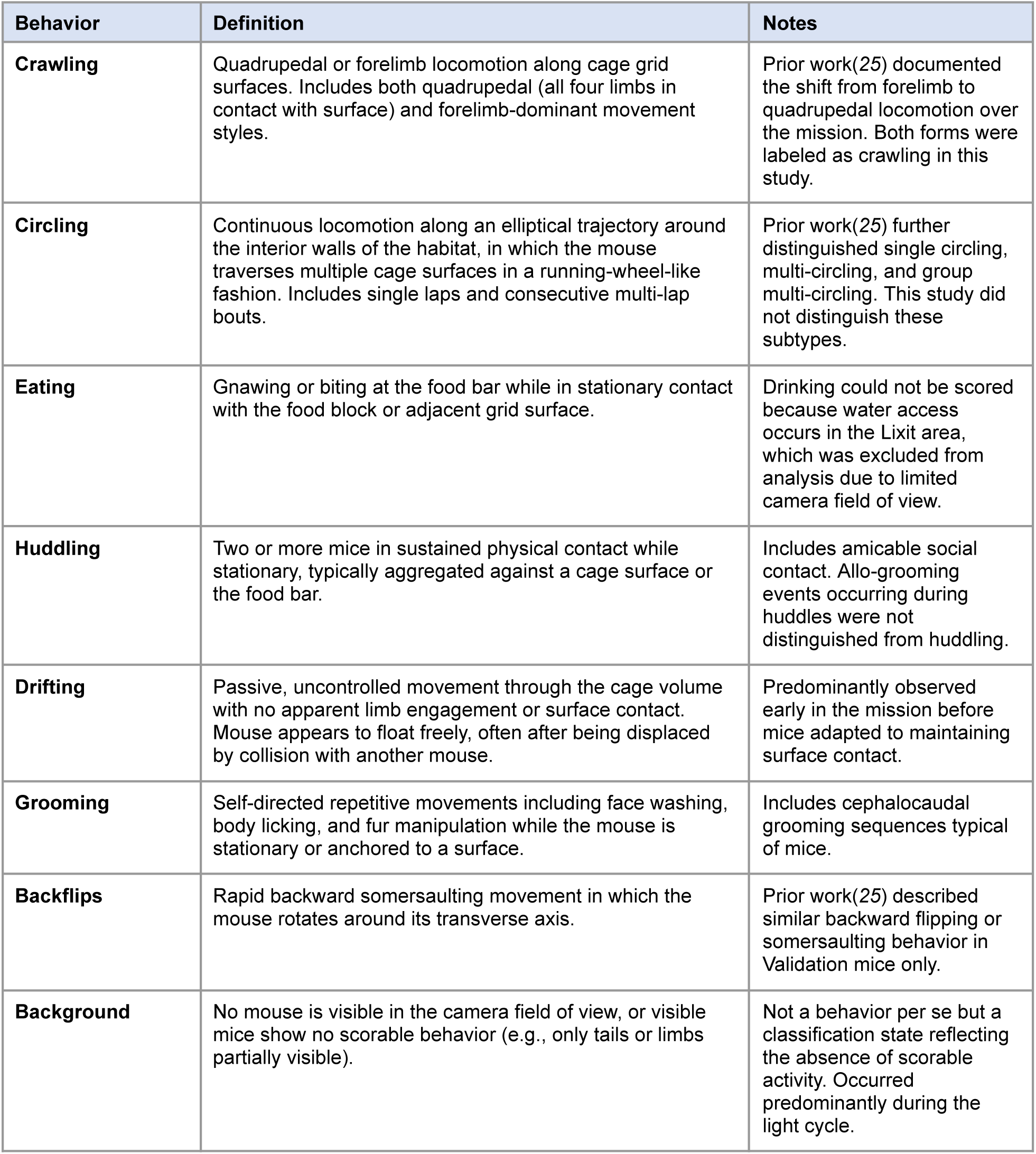
Operational definitions of the eight behavioral categories used for DeepEthogram labeling. Three behaviorists independently labeled *n* = 611,045 behavioral event-frames across 411,194 video frames from *n* = 66 videos. Behavioral criteria were adapted from prior work(*25*) and refined for reproducibility among labelers. Multiple behaviors could be labeled within a single frame because DeepEthogram supports multi-label classification.

